# Switching Logistic Maps to Design Cycling Approaches Against Antimicrobial Resistance

**DOI:** 10.1101/2020.03.17.995928

**Authors:** E.A. Hernandez-Vargas, C. Parra-Rojas, S. Olaru

## Abstract

Antimicrobial resistance is a major threat to global health and food security today. Scheduling cycling therapies by targeting phenotypic states associated to specific mutations can help us to eradicate pathogenic variants in chronic infections. In this paper, we introduce a logistic switching model in order to abstract mutation networks of collateral resistance. We found particular conditions for which unstable zero-equilibrium of the logistic maps can be stabilized through a switching signal. That is, persistent populations can be eradicated through tailored switching regimens.

Starting from an optimal-control formulation, the switching policies show their potential in the stabilization of the zero-equilibrium for dynamics governed by logistic maps. However, employing such switching strategies, deserve a specific characterization in terms of limit behaviour. Ultimately, we use evolutionary and control algorithms to find either optimal and sub-optimal switching policies. Simulations results show the applicability of Parrondo’s Paradox to design cycling therapies against drug resistance.

## 1 INTRODUCTION

Throughout history, we have witnessed alarming high death tolls derived from infectious diseases around the globe. Antimicrobials such as antibiotics and antivirals are powerful weapons to fight against infections. However, the misuse and overuse of drugs have led to *drug resistance*, which can be roughly defined as the ability of a microorganism to replicate in the presence of a drug [4]. Truly, during the course of an infection, pathogens can evolve genetically to generate resistance to a given drug. In other infections scenarios, the host can also be infected by different pathogenic variants deriving in a complex therapeutic challenge.

The World Health Organization (WHO) has reported that antimicrobial resistance (AMR) is a large-scale health problem worldwide [39]. This has been clearly exposed by HIV resistance to antiretrovitals [40] or the growing list of bacteria that are becoming harder to treat due to antibiotics becoming less effective e.g. pneumococcus, staphylococcus, aureus, pseudomonas aeruginosa among others [39].

Resistance can be developed by horizontal gene transfer of resistance encoding genes or mutations that derive the resistance phenotype to the population [6]. An important term is the *mutation rate*, which refers to the amount of genetic errors that accumulates per generation [15]. For example, mutation rates are about 10^−8^ to 10^−6^ substitutions per nucleotide per cell infection (s/n/c) for DNA viruses and from 10^−6^ to 10^−4^ s/n/c for RNA viruses [35]. Mutation rates in higher eukaryotes are roughly 0.003 mutations per genome per cell generation [14]. For different antibiotics, bacterial mutation rates oscillate between 3 × 10^−8^ to 5 × 10^−9^ per cell per generation [26].

Resistance can decrease the fitness of a pathogens, known as the biological cost of resistance, but in some cases, fitness can also be increased [3]. In this context, pathogens such as bacteria, viruses and fungus are mainly described respect to a standard wild type (most fit) in terms of phenotypes and genotypes [13]. While the *phenotype* attributes observable properties of the population, *genotype* refers to the genetic constitution. The scenario is multi-complex in the case of antibiotic resistance, for instance, mutations in different genes can produce similar antibiotic resistance phenotypes [5].

Mathematical modeling of infectious diseases has been developed at different scales [21]. Between-hosts models have helped to propose new vaccination strategies or support public health strategies [34, 16, 32]. On the other hand, for within-host infection, mathematical modeling has been used to capture the dynamics of different infectious diseases inside the host to understand the interaction of the pathogen and the immune system, as well as scheduling of therapies [2, 22, 11, 9, 33]. Most mathematical models to represent microbes dynamics are shown to be based on variations of the classical Verhulst logistic growth equation. For instance, the logistic model has served as a key mathematical tool in to represent the growth of tumors [1] and microbes [24]. The logistic model considers a stable population would consequently have a saturation level, known as the carrying capacity and forms a numerical upper bound on the growth size [38].

While theoretical approaches to mitigate drug resistance have been mainly developed at between-host level [37, 8], too little has been directed to investigate within-host strategies against antimicrobial resistance [23]. However, previous within-host control strategies are only developed for switched linear systems [21].

Here, for a any mutation network (illustrative example in Fig.1), we introduce a logistic switching map to capture the drug resistance dynamics of bacteria. This model is instrumental to design control strategies to minimize the ability of a bacterial sub-population to survive a drug concentration is known as *persistence*. However, designing switching strategies in dynamical systems is not trivial in the context of the so-called Parrondo’s Paradox [10, 27, 28], that is two losing games can be combined in a determined order to obtain a winning game [19]. Previous numerical simulations of switching logistic maps [28] have shown that switching decisions may follow either the “undesirable + undesirable = desirable” or the “chaotic+chaotic = order” dynamics.

**Fig. 1.**
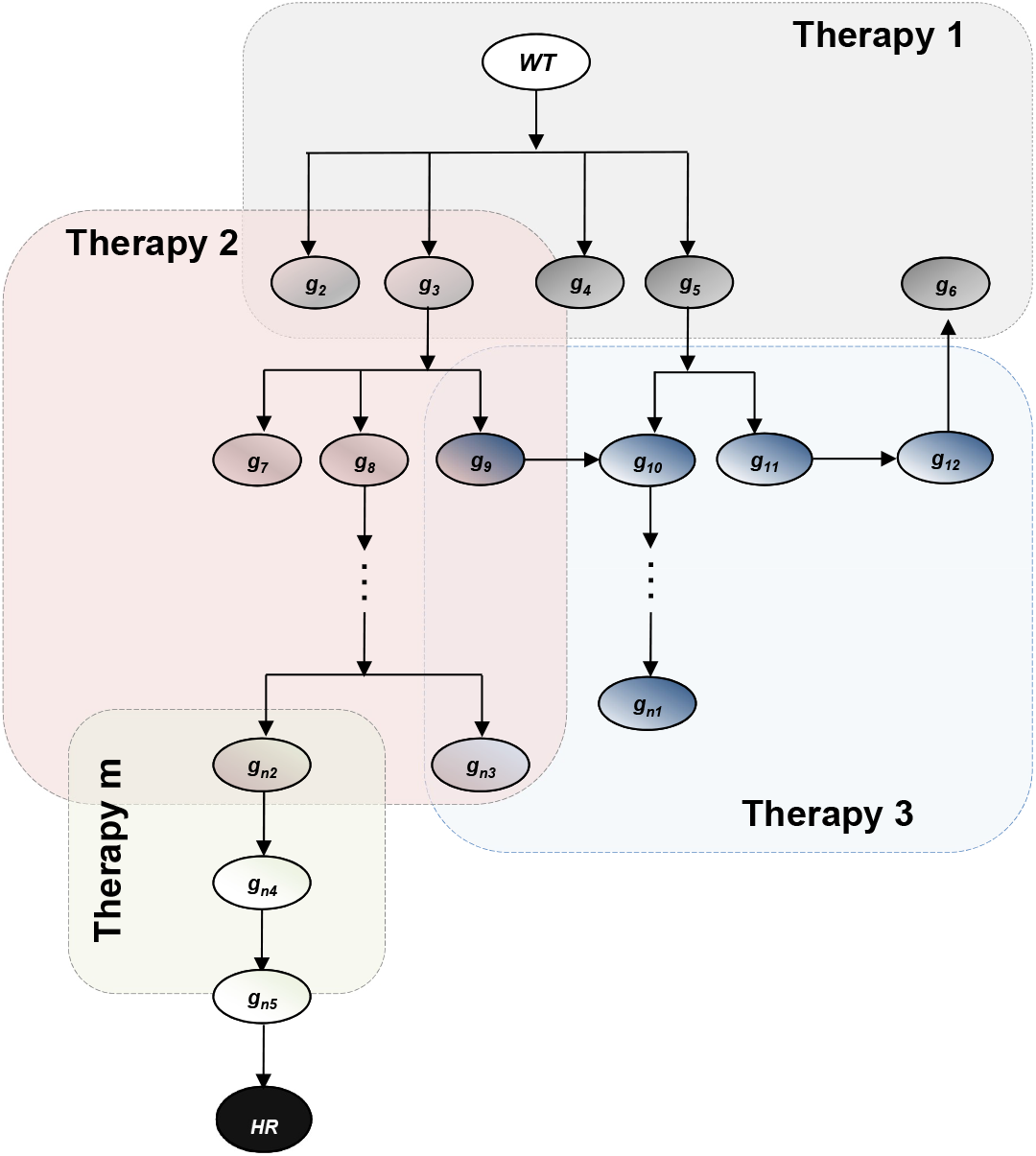
Illustrative mutation tree for *n* variants and *m* therapies. WT represents the wild type. Different variants would mutate the effect of a given therapy which is indicated with color squares. After a sequence of mutations, it is hypothesized the appearance of a highly resistant variant (black circle HR), meaning that it is resistant to both therapies.

Next sections, the mathematical abstraction of antimicrobial resistance dynamics is formulated in the form of non-linear switched systems. Consequently, we employ evolutionary and control algorithms to find sub-optimal switching policies. Simulations results show the applicability of Parrondo’s Paradox to design cycling therapies against drug resistance.

## 2 Logistic Switching Maps

Throughout, ℝ is the field of real number, ℝ^*n*^ stands for the vector space of all *n*-tuples of real numbers. ℝ^*n*×*n*^ is the space of *n* × *n* matrices with real entries. ℕ denotes the set of natural numbers. For *x* in ℝ^*n*^, *x*_*i*_ denotes the *i*^*th*^ component of *x*, and the notation *x* ⪰ 0 means that *x*_*i*_ ≥ 0 for 1 ≤ *i* ≤ *n*. 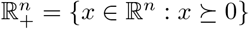 denotes the non-negative orthant in ℝ^*n*^. The transpose is represented as *A′*.

### Definition 1 Well-defined

A switching signal *σ*(·) is said to be *well-defined* on any interval [*t*_*k*_, *t*_*k*+1_), if it is defined in [*t*_*k*_, *t*_*k*+1_), and for all *t* ∈ [*t*_*k*_, *t*_*k*+1_).

### Definition 2 Well-posed

A switched system is said to be *well-posed* at *x*_0_ over any interval [*t*_*k*_, *t*_*k*+1_) if the switched system admits a unique solution via the well-defined switching signal *σ*(·) over interval [*t*_*k*_, *t*_*k*+1_).

Consider now the well-posed switched non-linear autonomous system described as follows:

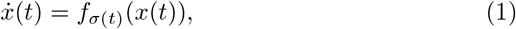

where 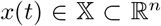 is the system state at time *t*, and the state space 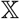 is closed. The initial condition at time *t*_0_ = 0 is *x*(0) = *x*_0_. *σ*(*t*)∈ℕ_*q*_ is the well-defined switching sequence that selects a transition function *f*_*i*_ ∈ ℝ^*n*×*n*^ for *i* = 1, 2,…, *q*, where *q* is a positive integer representing the number possible subsystems. Due to the applicability to biological systems, the set of non-linear functions represented by *f*_*i*_(*x*(*t*)) are positive, which is defined next.

### Definition 3 Positivity

For any initial condition 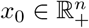 The set of continuous-time nonlinear sub-systems *f*_*i*_(*x*(*t*)) is called positive if 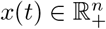 for all *t* ≥ 0

For a fix horizon of time *t*_*f*_ there is a finite length of horizon *t*_*f*_ − *t*_0_ which can be divided into *N*_*q*_ intervals. Thus, there is a well-defined switching path

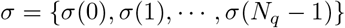

defined on the set ℤ_*N*−1_, which has a finite number of jump instants in any finite length sub-interval of 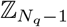. Any jump instant 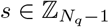 is said to be a switching time [18]. Thus, a switching time *s* satisfies *σ*(*s*) ≠ *σ*(*s* + 1). Let *s*_1_, *s*_2_, ⋯, *s*_*l*_ be the ordered of switching times in 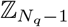 with

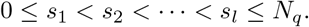

The ordered sequence 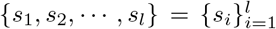, is defined as the *switching sequence* of *σ* on 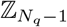, which implies that there are *l* jump instants. The *i* element of the path *σ*, then

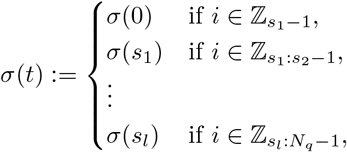

which means that *σ* is such that

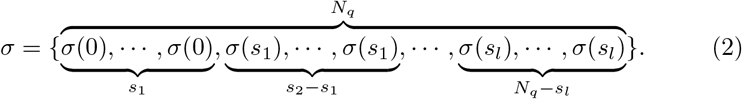

Now, let *h*_*i*_ = *s*_*i*+1_ − *s*_*i*_ for *i* = 0, ⋯, *l* (consider *s*_0_ ≔ 0), then the sequence {(*σ*(0), *h*_0_), ⋯, (*σ*(*s*_*l*_), *h*_*l*_)} is said to be the *switching duration sequence* of *σ* on 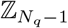. It is clear that the switching duration sequence is uniquely determined by the switching sequence, and vice versa.

### Remark 1

Different initial conditions will correspond to distinct switching paths and hence different switching sequences. Thus, the switching sequence depends heavily on the initial state [18].

From literature in switched systems [27], we know that a switched system is stable if all individual subsystems are stable and the switching signal between them is sufficiently slow. Next, we bring the definition of dwell-time.

### Definition 4 Dwell-time

The switching times *t*_*i*_ satisfy the inequality *t*_*i*+1_ − *t*_*i*_ ≥ *τ*_*d*_ for all *i*, where *τ*_*d*_ ≥ is the dwell-time.

### Problem 1

In this work, *σ*(·) is considered as the only manipulable control signal to the system (1). The primary goal, if possible, would be to design a switching path to make the origin of the system stable (1). However, for biological reasons of the application described in next sections, we relax the problem to the design of a sub-optimal switching path *σ*(·) that yields the closest behavior to satisfy a cost functional 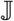 in a fix horizon of time (*t*_*f*_).

### 2.1 Modeling Antimicrobial Resistance as Logistic Maps

To abstract the dynamics of the different genotypes during drug therapy, the following switched logistic model is described:

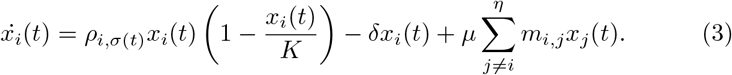

The population of different *η* pathogenic variants are represented with *x*_*i*_. *ρ*_*i*,*σ*(*t*)_ is the proliferation rate of the variant *i* under the treatment regiment *σ*(*t*) ∈ {1, 2, 3,…, *N*} which can be changed at any time *t*. *N* is the total number of possible drug therapies that can be administered. The maximum carrying capacity is *K*. *δ* is the clearance of the variant *x*_*i*_.

The mutation rate is represented by *μ*. The genetic connections between genotypes is represented by *m*_*i*,*j*_ ∈ {0, 1}, that is, *m*_*i*,*j*_ = 1 if and only if it is possible for genotype *j* to mutate into genotype *i*. An illustrative example to represent the mutation network for nine variant and two therapies is presented in Figure 1. However, more complex mutation tress can be considered. The switched model (3) is based on following assumptions:

#### Assumption 1 Negligible effect of the immune system

*For a complete picture of a within-host infection and the corresponding selective pressure that could derive into a new phenotype, the immune responses would need to be modelled. However, this would result in a complex model with many parameters to fit and still difficult to represent the reality [22, 12]. As we focus to chronic infections that could not been cleared by the immune system, the main pressure and clearance of pathogenic variants would be directed by the drug. This assumption allows to simplify the model to being essentially a logistic map of the pathogen dynamics.*

#### Assumption 2 Logistic deterministic dynamics

*Logistic equations is how most of bacterial infections are modelled [24, 38]. A main interest in this study is to compare bacterial growth of different sub-populations with control strategies under distinct infection scenarios, thus logistic are constructed based on ODEs. While this will simplify the control design, it is also a strong assumption, as it will be difficult to evaluate numerically if a control strategy derived in eradication. The alternative would be a stochastic model, which is in fact a very attractive long-term problem*.

#### Assumption 3 Pathogen clearance independent of therapy and mutant

*Pathogen clearance rate (parameter δ) could depend on one or more of the treatment regimes or the variant genetics. For the case of viral infections, antivirals can only inhibit the viral replication inside of cells [21]. For the case of bacterial infections, penicillin-based antibiotics can kill directly bacteria while bacteriostatic inhibits bacterial proliferation [25]. Mathematically speaking, the drug effect is effective if the pathogenic clearance is bigger than its proliferation (δ* > *ρ*_*i*_*). For simplicity to design switching policies, this is considered constant for all the variants.*

#### Assumption 4 Resources are not a limited factor

*During the course of an infection, the amount of sources (e.g. nutrients, cells to infect, etc.) are usually in excess. Thus, we assume the same current capacity (parameter K) for the different variant during the course of the therapy*.

#### Assumption 5 Mutation rate independent of treatment and strain

*There is the possibility of dependence of mutation rate (parameter μ)) on the replication rate. Thus, there is a relationship between genetic strain, treatment, and mutation rate. For simplicity in the design of control strategies, it is assumed that the mutation rate between species with the same genetic distance is constant*.

#### Assumption 6 Growth rates based on therapy

*The most adequate framework to represent the drug effects on a disease is with the corresponding PK/PD dynamics (pharmacokinetics and pharmadynamics). To quantify bacterial resistance, the minimum inhibitory concentration (MIC) is used to measure the lowest concentration of an antibiotic to prevent bacterial replication [4]. Previous control engineering works [20] formulated the scheduling of drugs in an impulsive framework. While this is an excellent framework for single therapy, the complexity increase when different therapies are considered. Thus, it is assumed that a corresponding therapy affect instantaneously the pathogen, which is a reasonable assumption for chronic diseases due to the long-term time scales*.

## 3 Stability analysis without Switching

Let us take the system (3) and focus on the case of a single therapy. Without loss of generality, we simplify the notation by taking *ρ*_*i*,*σ*(*t*)_ → *ρ*_*i*_, and normalizing the bacterial populations by the carrying capacity of the system, *x*_*i*_/*K* → *x*_*i*_, so that

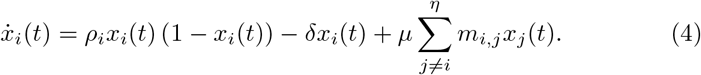

The strains that mutate act as a *source* to one or more strains, whereas each strain can have at most one source, with the case of zero sources corresponding to the wildtype. In other words, we do not take into account the case when two or more different strains can yield exactly the same genotype upon mutation, since this is unlikely in a real-life scenario. At the same time, and for the same reason, all *backwards* mutations are forbidden, so that if *m*_*j*,*i*_ ≠ 0 then necessarily *m*_*i*,*j*_ = 0. More generally, there is no way of coming back to strain *i* starting from strain *i*; that is, once a mutation from strain *i* to strain *j* occurs, there exists no possible sequence of *L* mutations *x*_*j*1_ → *x*_*j*2_ → … *x*_*jL*_ → *x*_*i*_ for any length *L*.

The equilibrium points of the system with *n* strains, which we denote by 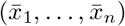, can be found recursively by solving for 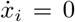 in Eq. (4), and have the form

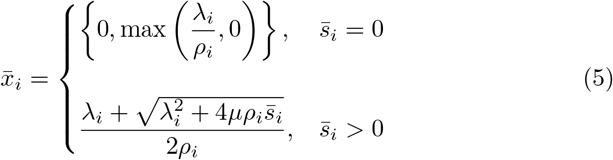

Here, we have denoted by 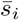 the equilibrium value of the strain that acts as a source of *x*_*i*_; this means that, if *x*_*i*_ results from the mutation of a certain *x*_*k*_, then 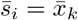. At the same time,

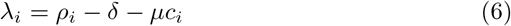

where *c*_*i*_ is the number of *children* of strain *i*, *i.e.*, the number of genotypes that result from one mutation of genotype *i*—we note that *c*_*i*_ = −*m*_*i*,*i*_.

### Theorem 1

λ_*i*_ > 0 *ensures the existence of the equilibrium point with* 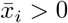 *and* 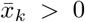 *for all x*_*k*_ *that can be reached starting from x*_*i*_, *with* 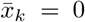 *otherwise. This point is linearly stable if, additionally*, λ_*k*_ < 0 *for all other k*.

*Proof* The Jacobian of the system evaluated at a given equilibrium point can be decomposed as *J* = *D* + *μM*_1_, where *M*_1_ contains the off-diagonal elements of *M*, and thus only encodes the mutations between strains. The entries of *D* are given by

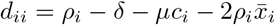

The properties derived above for the constraints on the mutation patterns imply that *M*_1_ can be thought of as the adjacency matrix of a directed acyclic graph (DAG) [29]; therefore, there exists a topological ordering of the strains that renders *M*_1_ strictly upper triangular [30]. In other words, we can relabel the strains so that for every mutation from *x*_*i*_ to *x*_*j*_, *i* comes before *j* in the ordering. With this, the Jacobian becomes upper triangular, and its eigenvalues will be given by the entries of *D*. At the origin, the *i*-th eigenvalue will correspond to λ_*i*_ from Eq. (6). On the other hand, the eigenvalues associated with non-zero coordinates are given by

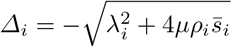

and they are always negative. As a consequence, whenever a strain *x*_*i*_ is able to persist by itself—which translates into λ_*i*_ > 0—then all of its children will also persist, as seen from Eq. (5), even if they would be eradicated in isolation. If all other strains in the system are unable to persist in isolation, the eigenvalues associated to the equilibrium point where 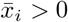, with 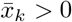 if *k* corresponds to a children of *i*—that is, 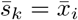—and 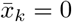 otherwise, will be given by

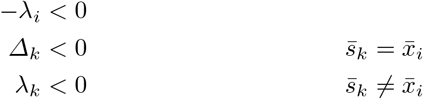

and the equilibrium is thus linearly stable.

### Theorem 2

*If we explicitly separate the zero and non-zero equilibrium coordinates—so that the i-th element of an equilibrium point is either 0 or* 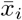—*then the set defined by*

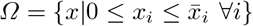

*is positively invariant with respect to the system described by* *Eq*. (4)*. That is, for any given initial condition x*_0_ ∈ *Ω when t* = 0, *the solution to* (4) *satisfies x*(*t*) ∈ *Ω* ∀*t* > 0 *[7]*.

*Proof* Let us recall that 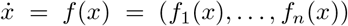, and consider the boundaries of *Ω*, given by

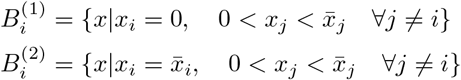

The invariance condition translates into 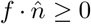 along the boundaries above, where 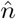 is a normal vector pointing towards *Ω*. We note that the normal vectors that correspond to each boundary are given by 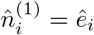 for 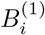 and 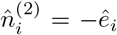 for 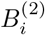, where 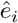 is the *i*-th unitary vector, 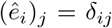, with *δ*_*ij*_ the Kronecker delta.

Choosing an initial condition *x*_0_ along 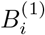, with 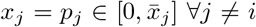 we have

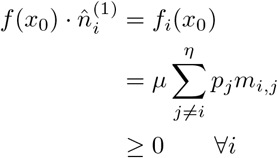

Similarly, for the case of 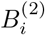, we choose an initial condition *x*_0_ with 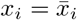 and 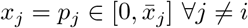; this yields

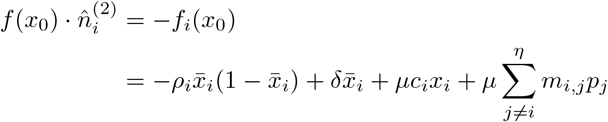

Now, we use the fact that 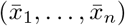 is an equilibrium point of the system, so that

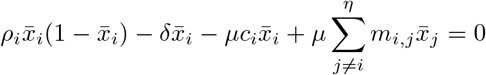

and

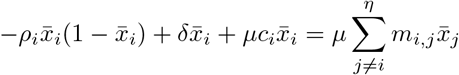

Hence,

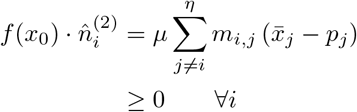

since 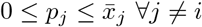.

Therefore, *Ω* is positively invariant with respect to the system (4).

### Corollary 1

*The origin of the system is globally asymptotically stable when*

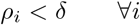

*Proof* We have seen above that the origin is a linearly stable equilibrium point of (4) when *ρ*_*i*_ < *δ* + *μc*_*i*_ ∀*i* simultaneously. In order to establish the global stability of the origin, let us choose a Lyapunov function

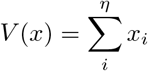

We note that *V* ≥ 0 ∀*x* ∈ *Ω*, and *V* = 0 iff *x* = 0. Now,

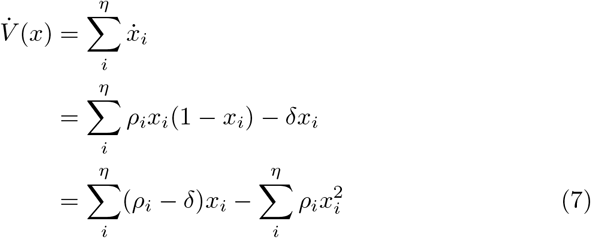

From the expression above, *ρ*_*i*_ < *δ* ∀*i* ensures that 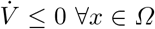, and 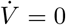 iff *x* = 0. Therefore, the origin is globally asymptotically stable under this condition.

The condition above arises trivially from considering all strains as independent of one another, in absence of mutations, and it is more restrictive than the condition found previously, since mutations have a stabilizing effect on the origin. However, Eq. (7) can be employed as a basis to establish a switching policy in order to ensure the eradication of the pathogen in the case with mutations under a set of different available therapies.

## 4 Switching therapies

Based on the general model (3), the problem description can then be broadly described as the design of a switching policy for the gradual eradication of pathogenic strains. To formally state the problem we need the following definitions:

### Definition 5

A switching policy is the cycling choice between therapies *σ* with a period *T*_*σ*_, which can be represented by a piecewise constant function *σ*(*t*) : ℝ → {1, 2,…, *N*}. *σ*(*t*) remains constant for all *t* ∈ [*kT*_*σ*_, (*k* + 1)*T*_*σ*_)

### Definition 6

A periodic cycling policy is the periodic switching between therapies *i* with regular intervals *T*_*i*_.

For example, for the case of two therapies with periods *T*_1_ and *T*_2_, we have *σ*(*T*_1_) = 1 and *σ*(*T*_2_) = 2, then the periodic cycling policy sequence is

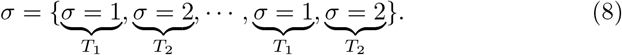

Next, we describe the design of switching policies of the system (3) in discrete-time version. To this end, we will divide the analysis in two parts. The first is assuming the variants do not mutate that is *μ* = 0, this is still an interesting biological scenario as the main goal is to eradicate different variants of a pathogenic microorganism. The second part will consider mutation, as *μ* ≠ 0 for the case of not backward mutations, as from the evolution point of view, it is very difficult that this type of mutations occur.

### 4.1 Switching Policies without mutation *μ* = 0

Assuming no mutations (*μ* = 0) we re-parameterized the model (3) for a fixed period of treatment (*σ*(*t*) = *σ*) as follows:

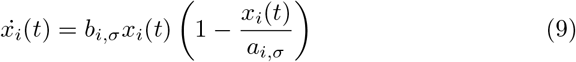

where *b*_*i*,*σ*_ = *ρ*_*i*,*σ*_ − *δ* and *a*_*i*,*σ*_ = *Kb*_*i*,*σ*_/*ρ*_*i*,*σ*_. Equation (9) has the conventional logistic form which can be solved analytically, thus the recursive form of (3) is as follows:

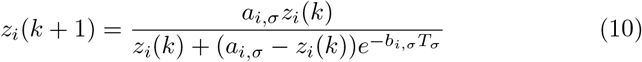

where *T*_*σ*_ is the period where the therapy *σ* is provided.

#### Theorem 3

*Given the dynamics in (3) without mutation (μ* = 0*) and any periodic cycle (T*_*σ*_*) of the different therapies (σ* = 1,…, *N), the different variants will go to eradication independently if for a given treatment the origin is unstable if and only if the following condition is satisfied*

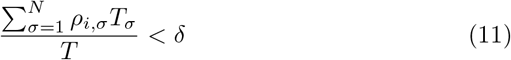

*where* 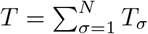.

*Proof* Considering the initial condition *z*_0_ = *z*(0) = 0 and the first two cycle of therapies *σ* = 1, 2 with their respective period *T*_*σ*=1,2_, by construction, we have the the Jacobian matrix *J*_*σ*=1,2_ at the origin as follows:

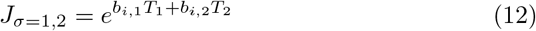

In similar vein we can generalize the Jacobian *J*_*σ*=1,..,*N*_ in the origin as follows,

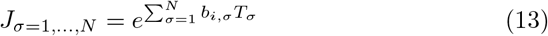

as *b*_*i*,*σ*_ = *ρ*_*i*,*σ*_ − *δ* and and we are interested in the stability of the origin, we check the exponential term to be less than 1, then

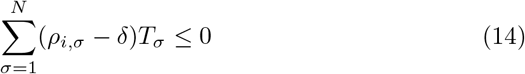

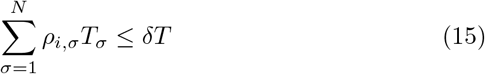

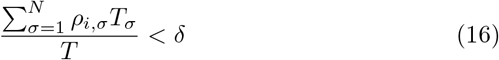

#### Remark 2

The condition (11) can additionally serve to design the duration of the therapies (*T*_*σ*_) to eradicate all different variants during the course of an infection if the following inequality system has a solution

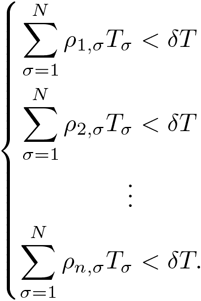

This can be formulated as a feasible solution to a linear programming problem. While this could provide a sufficient condition for a possible success of a cycling therapy, the unfeasible solution does not necessarily imply the failure of a switching trajectory to stabilize the origin.

#### Corollary 2

*For a cycle of the therapies with equal time duration, the stability condition (11) can be further reduced as*

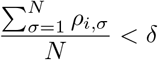

*Proof* Following the Theorem 3, we have that the period of each therapy is equal, that is *T*_1_ = *T*_2_ = … = *T*_*N*_. Then, 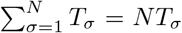. Therefore *T*_*σ*_ can be deleted from numerator and denominator in conditional (11).

#### Remark 3

The corollary 2 points out that in order to eradicate a variant *i* with cycling therapies, the average expansion of the variant *i* through the cycling should be less than its contraction.

### 4.2 With Mutation *μ* ≠ 0

For the case with mutation (*μ* ≠ 0), the condition (11) is difficult to proof. Model Predictive Control (MPC) appears to be suitable for a suboptimal application to the biomedical application, due to its robustness to disturbances, model uncertainties and the capability of handling constraints [21–23, 41]. Thus, selection of therapies based on MPC will be employed on similar reasoning as in our previous work based on switched linear systems [23]. In summary, if the total population size is small enough during a finite time of treatment, then there is a significant probability that the population becomes zero. Therefore, we consider the cost

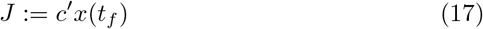

where *c* is the column vector with all ones, and *t*_*f*_ is an appropriate final time. This cost should be minimized under the action of the switching rule. MPC problem can be formulated as in [17]. Based on measurements obtained at the step *k*, the controller predicts the future dynamic behavior of the system over a prediction horizon *T_p_* and computes an open-loop optimal control problem with control horizon *T*_*c*_, to predict the future input for the system. The problem is written as follows:

#### Problem 2

The internal variables in the controller by a bar 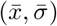 is denoted, where 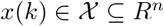 and 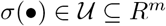. Find

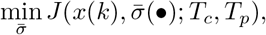

with

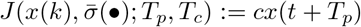

subject to:

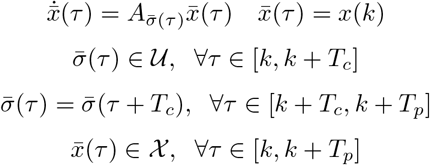

The MPC involves a nonlinear and complex optimization problem, the global optimization algorithm known as differential evolution [36] is considered here.

## 5 Numerical Simulations

In similar way to [23], simple illustrative examples are considered based on 4 genetic variants and two different therapies. Thus, *η* = 4, and 2 drug therapies, *N* = 2. The strain *i* = 1 is considered the Wild Type which is susceptible to most of drugs. The Genotype 1 (*G*_1_) is resistant to therapy 1 but it is susceptible to therapy 2. The Genotype 2 (*G*_2_) is resistant to therapy 2, but it is susceptible to therapy 1. The Highly Resistant Genotype (*R*_*G*_) is a genotype with low replication rate, but it is resistant to all drug therapies.

Pathogen clearance rate is fixed, *δ* = 0.24 *day*^−1^, which corresponds to a half life of slightly less than 3 days [31]. Typical mutation rates are of the order of *μ* = 10^−4^. The following mutation matrix is considered:

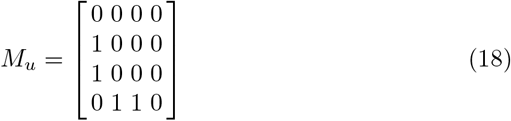

Three different scenarios are proposed as in the Table 1. The first scenario, the most ideal case, describes a stabilizable switched system with a complete symmetry between *G*_1_ and *G*2, in the sense that therapy 1 inhibits *G*_2_ with the same intensity that therapy 2 inhibits *G*_1_. In practice, a small difference in relative replication ability is expected. The second scenario shows a system that can not be stabilized with a complete symmetry between *G*_1_ and *G*_2_ (similar to Case 1) but there is a resistant genotype that can not be stabilize under switching. The third scenario is a switched system that can be stabilize under switching and it has completely asymmetry for replication rates in *G*_1_ and *G*_2_. In general, the finite set of possible control values causes problems for many control techniques. Nevertheless in the cause of MPC, having a finite set of options may be an advantage in making the optimization easier to solve.

**Table 1.** Illustrative simulations scenarios for drug Resistance based on Equation (3)

| Scenario | Therapy | WT($x_1$) 1 | $G_1$ ($x_2$) | $G_2$ ($x_3$) | $R_G$ ($x_4$) |
| --- | --- | --- | --- | --- | --- |
| 1 | 1 | $\rho_{1,1} = 0.05$ | $\rho_{2,1} = 0.27$ | $\rho_{3,1} = 0.05$ | $\rho_{4,1} = 0.20$ |
| | 2 | $\rho_{2,1} = 0.05$ | $\rho_{1,2} = 0.05$ | $\rho_{3,2} = 0.27$ | $\rho_{4,2} = 0.20$ |
| 2 | 1 | $\rho_{1,1} = 0.01$ | $\rho_{2,1} = 0.3$ | $\rho_{3,1} = 0.1$ | $\rho_{4,1} = 0.3$ |
| | 2 | $\rho_{2,1} = 0.01$ | $\rho_{1,2} = 0.1$ | $\rho_{3,2} = 0.3$ | $\rho_{4,2} = 0.3$ |
| 3 | 1 | $\rho_{1,1} = 0.01$ | $\rho_{2,1} = 0.26$ | $\rho_{3,1} = 0.05$ | $\rho_{4,1} = 0.25$ |
| | 2 | $\rho_{2,1} = 0.01$ | $\rho_{1,2} = 0.15$ | $\rho_{3,2} = 0.38$ | $\rho_{4,2} = 0.24$ |

To compute optimal switching trajectories, we consider a “brute force”, which calculates all possible combinations of the therapies (*N*_*c*_ = *t*_*f*_/*T*_*c*_) and then finding the minimum for the cost function (17). However, this approach is computational unfeasible for checking *N*_*c*_ > 15. Thus, to check larger treatment combination we consider the differential evolution algorithm [36]. Furthermore we consider also a MPC strategy as discussed in Problem 2.

Table 2 illustrates treatment scenarios of 200 days (*t*_*f*_) with decision time to switch therapy of 20 days (*T*_*c*_), that is *N*_*c*_ = 10, 2^10^ combinations. Based on the column of “brute force” results in the Table 2, we can conclude that the DE algorithm can find optimal trajectories. While we can not guarantee DE algorithm will find the optimal solution for all any example, it might find solutions very close to the optimal. In similar fashion, the MPC was able to find optimal switching trajectories in case 1 and 2 but not in case 3. However, we can observe it find solution very close to the optimal.

**Table 2.** Total pathogen load at the end of treatment of 200 days with a possibility to switch every 20 days.

| Scenario | Optimal (Brute Force) | Optimal (DE Algorithm) | MPC |
| --- | --- | --- | --- |
| 1 | $2.3615 \times 10^{-5}$ | $2.3615 \times 10^{-5}$ | $2.3615 \times 10^{-5}$ |
| 2 | $1.8851 \times 10^3$ | $1.8851 \times 10^3$ | $1.8851 \times 10^3$ |
| 3 | 0.1040 | 0.1040 | 0.1141 |

The advantage of MPC would be the computational time, thus we could explore short decision times. Thus, in Fig. 2 we illustrate the performance of MPC for a horizon of 500 days (*t*_*f*_) with decision time to switch therapy of 5 days (*T*_*c*_), that is *N*_*c*_ = 100, 2^100^ combinations. This scenarios would not be computational feasible for a “brute force”. The upper panel in Fig. 2 shows how the MPC strategy can decrease the pathogen load of the different genotypes. The lower panel in Fig. 2 presents the switching trajectory, which is not having any intuitive patron, thus highlighting the potential of MPC strategy.

**Fig. 2.**
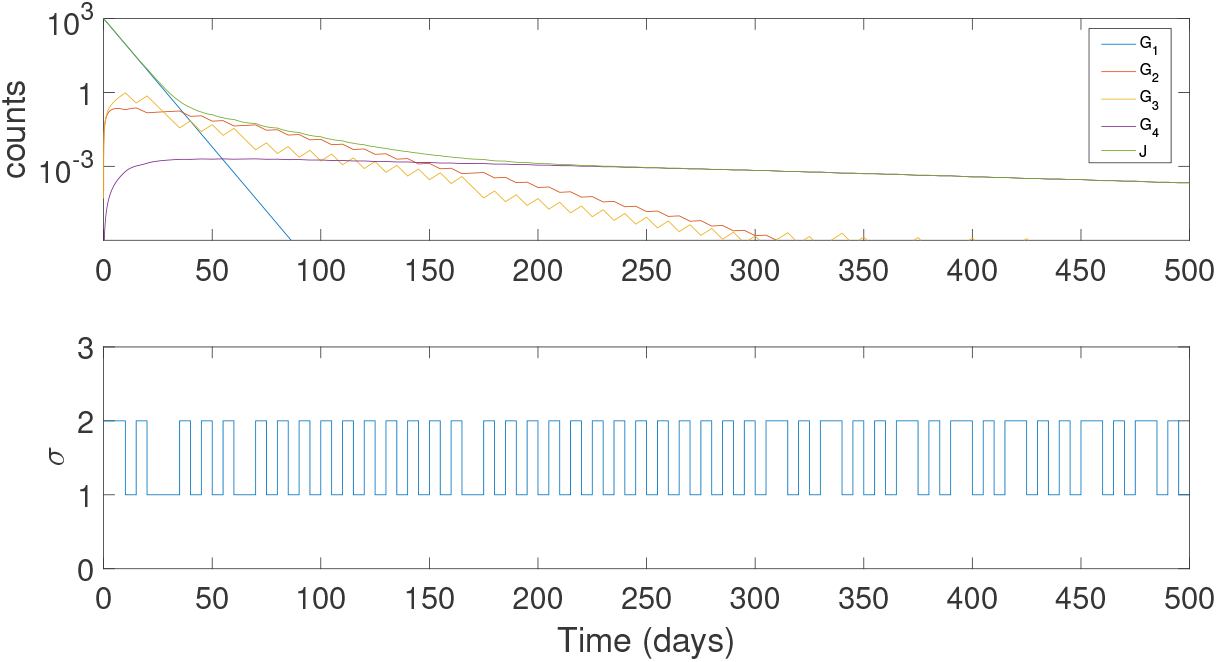
Numerical results for simulation scenario 3. This considers a simulation time of 500 days with a decision time of every 5 days. The wildtype is rapidly decreased. The genotypes 2 and 3 are gradually decreased by the switching between therapies. It can be observed that the switching rule is not having a clear patron. The resistant genotype is also decreased due to the appropriate switched between the therapies. Total pathogen load is represented in the cost function *J*.

## 6 Conclusions

This paper introduced a switching logistic model wit the potential to be the basis for scheduling antimicrobial to mitigate resistance. Considering the case of not treatment and mutation trees where backwards mutations are forbidden, we derived conditions of global stability For the case of therapies without mutations, conditions for which unstable zero-equilibrium of the logistic maps can be stabilized through a switching signal. For the case with mutations, computational switching strategies such as MPC were formulated. Numerical results highlighted that MPC strategy will perform very close to optima control while computational resources are largely decreased.

